# Graph Signal Processing on protein residue networks helps in studying its biophysical properties

**DOI:** 10.1101/2021.01.02.425090

**Authors:** Divyanshu Srivastava, Ganesh Bagler, Vibhor Kumar

## Abstract

Understanding the physical and chemical properties of proteins is vital, and many efforts have been made to study the emergent properties of the macro-molecules as a combination of long chains of amino acids. Here, we present a graph signal processing based approach to model the biophysical property of proteins. For each protein inter-residue proximity-based network is used as basis graph and the respective amino acid properties are used as node-signals. Signals on nodes are decomposed on network’s Laplacian eigenbasis using graph Fourier transformations. We found that the intensity in low-frequency components of graph signals of residue features could be used to model few biophysical properties of proteins. Specifically, using our approach, we could model protein folding-rate, globularity and fraction of alpha-helices and beta-sheets. Our approach also allows amalgamation of different types of chemical and graph theoretic properties of residue to be used together in a multi-variable regression model to predict biophysical properties.

## 1. Introduction

Proteins are the fundamental building blocks of a cell. The sequence of amino-acids (or residues) is stabilized into a native, functional three-dimensional state of the protein. Thus, the smaller building blocks of the protein emerge as functional only when they are arranged in a defined arrangement in a three-dimensional space [1]. It is a rather well-known fact that protein’s overall biophysical behaviour is governed by this peculiar order and residues arrangement. Therefore in addition to protein sequence information and secondary structure, researchers are also discovering new ways to utilize tertiary structure to model properties of proteins.

Multiple attempts have been made to model a protein molecule as network of residues as nodes connected with each other based on the distance between them in 3D structure of a protein [2, 3, 4]. Network properties of nodes (residues) based on graph theory such as centrality, betweenness and clustering coefficient have been used to predict biophysical properties using protein structures [3]. Another instance is community network analysis (CNA) which is used to study the dynamics of enzymes and protein/DNA (and/or RNA) complexes for understanding their allosteric mechanisms[5] [6]. Similarly, the folding rate of protein has also been modelled using network properties based on only graph-theoretic approach [7]. Though such approaches highlight the importance of residue network properties, however ignoring the biophysical properties of amino acids could lead to under-utilization of previously available information about the residues. Thus the question arises how to efficiently amalgamate different kinds of signals due to amino acid properties and 3D proximity neighbourhood information.

Here we propose a conceptually different approach using graph signal processing, which amalgamates residue properties and residue network structure to model proteins’ biophysical properties. Notably, we show how a protein’s structure information and residue’s biophysical properties influence folding rate of protein. Further, we show how different kinds of features of amino acid residues can be combined with structural information to apply regression models to predict 3 types of properties of protein.

## 2. Materials and Methods

Throughout this study, protein molecules are processed as graphs. A graph *G*(*V, E*) is a collection of *V* vertices and *E* edges. The entire framework from constructing these graphs to the processing’s mathematical foundations is described in this section.

### 2.1. Protein Network Models

#### 2.1.1. Residue Interaction Graph

A residue interaction graph (RIG) model of a protein is a simple network model of a protein. A simple RIG model for a protein is a graph, in which each vertex corresponds to a residue, and two residues are connected with an edge, if they lie in proximity in their native-state structures. Given the three-dimensional coordinates of each atom in a protein (available in the Protein Data Bank [8]), inter-residue Euclidean distances are calculated (Figure 1b). For consistency, the centre of a residue is considered to be its alpha carbon (*C_α_*) atom [9]. Two vertices *υ*_1_ and *υ*_2_, with their corresponding *C_α_* coordinates (*x*_1_, *y*_1_, *z*_1_) and (*x*_2_, *y*_2_, *z*_2_) are connected if the distance between the two is below or equal to a certain threshold *r_c_*. In general, we have

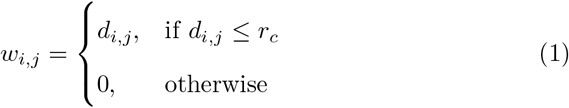

where *r_c_* is the cutoff distance, and 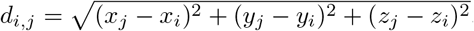. The cutoff *r_c_* is carefully chosen to consider the required forces of attractions which indeed keep the protein structure stable. A suitable and meaningful cutoff *r_c_* is usually chosen in the range 5-9 Å [4].

**Figure 1:**
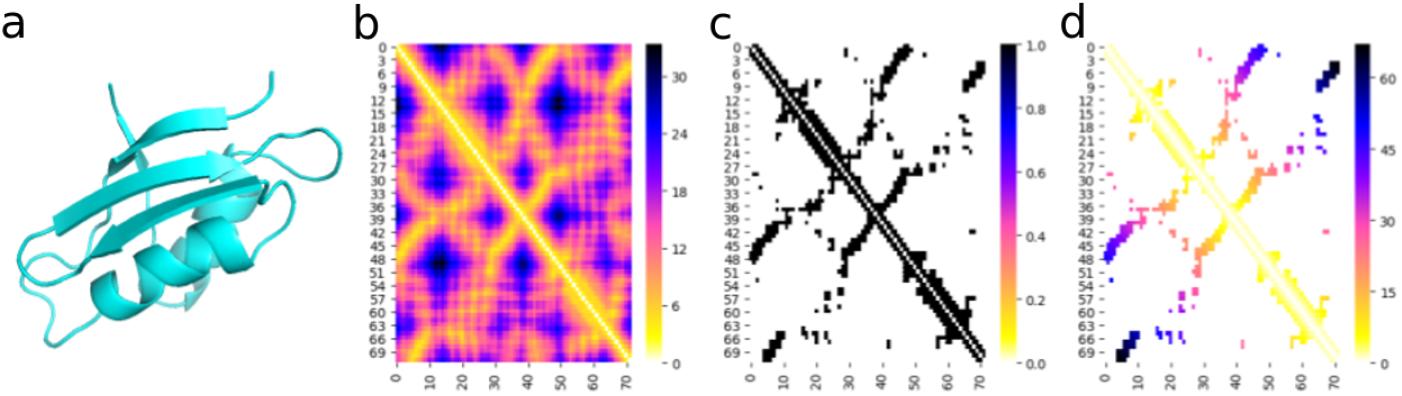
Construction of a weighted Residue Interaction Network for a protein (PDB ID : 2HQI). (a) A stable tertiary structure of a protein (b) the inter-residue Euclidean distances and all-vs-all contact map (c) The contact map is a heat map of the inter-residue distances of the protein, and is analogous to the network’s adjacency matrix. The distance cutoff (*r_c_*) is chosen as 8 Å in this example, and a binarized unweighted RIG model is created (d) Weights are added to the unweighted RIG model according to distance in sequence of protein, to obtain to weighted RIG model’s adjacency matrix.

A slight variant of the simple RIG model is the unweighted RIG model, which is a binarized version of the simple RIG model. The model’s weights are defined as

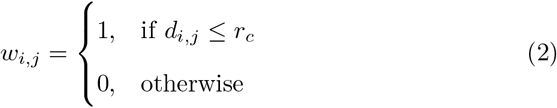

Figure 1c shows the adjacency matrix for unweighted RIG model of a human protein (PDB ID: 2HQI), where *r_c_* = 8 Å.

#### 2.1.2. Weighted RIG model

While the RIG model captures the overall three-dimensional structure of the protein to some extent, it fails to highlight the interactions among residues lying distant apart along the backbone of the protein sequence. Such long-range interactions are the ones majorly responsible for holding the structure intact and aids protein folding [10]. Long-range Interaction network have been studied previously in the context of protein folding kinetics [2]. Here, we use a modified model which includes both short and long-range interactions. It takes all edges of the RIG model into consideration and weights the edges such that long-distance edges have higher edge weights (Figure 1d). The weights are proportional to the distance between the residues along the backbone of the protein. Thus, long-range interactions are given more importance. This model is referred to as the Weighted RIG (*wRIG*) model. The edge weights are calculated as -

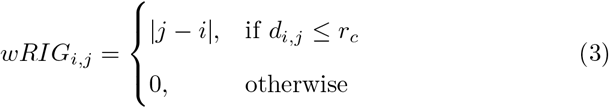

This model is shown to have produced better results as compared to other models in the later sections.

### 2.2. Spectral Graph Theory

Spectral graph theory is the study of the eigenvalues and eigenvectors of the matrices associated with the graph[11]. The (normalized) Laplacian matrix is often used for the purpose of graph signal processing, which is defined as

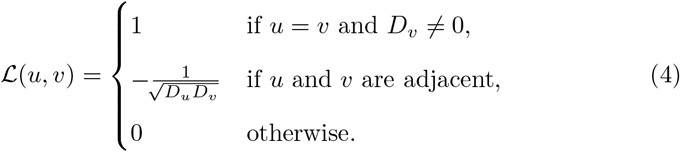

where *u, υ* are vertices in the network, and *D_i_* is the total degree of the *i^th^* vertex. The Laplacian is also a square matrix like the adjacency matrix. The Laplacian matrix is a positive semi-definite matrix, hence it has a complete set of real and orthonormal eigenvectors. Moreover, since the matrix is symmetric with real values, it is diagonalizable. The eigendecomposition of 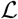 is given as 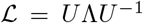, where *U* and Λ are the eigen basis matrix and a diagonal matrix of eigenvalues of 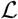 respectively. Now, since *U* is the matrix of orthonormal eigenvectors, we have, 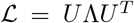. The corresponding eigenvalues are non-negative. The eigenvectors are denoted by {**u***_l_*}_*l*=0,1,…*N*−1_. Zero occurs as an eigenvalue in multiplicity of the number of connected components of the graph. The eigenvectors of L are denoted by 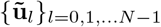. The eigenvalues _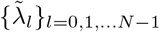_ of the normalized graph Laplacian matrix satisfy the condition 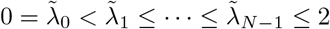.

### 2.3. Graph Signal Processing

For a graph *G*(*V, E*) with *V* vertices and *E* edges, a graph signal is represented as a vector *f* ∈ ℝ_*V*_, such that the *i^th^* component of the vector represents the value of the signal at vertex *υ_i_*. A graph Fourier basis is obtained through the spectral decomposition of its Laplacian Matrix [12]. For a signal *f* defined on the vertices of a graph, its Graph Fourier Transform (GFT) is defined as

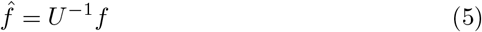

Where *U*^−1^ is the *Graph Fourier transform matrix*. Since *U* is the matrix of orthonormal eigenvectors, *U*^−1^ = *U ^T^*. The values of 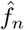 of the signal’s graph Fourier transform characterizes the frequency content of the signal as a projection on the eigenvector **u***_n_* [13].

The inverse graph Fourier transform (I-GFT) is given by

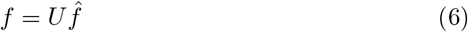

It constructs the original back from its frequency components. According to GSP theory, the transformed signal 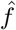 is analogous to the frequency domain transformation [13]. Since we use Laplacian of graph, the components towards the smaller eigenvalues represent low frequency while the high eigenvalue components represent higher frequency [13]. The higher frequency components are often considered to be dominated by noise.

In this study, various biophysical attributes of the residues, like their molecular weight, hydrophobicity etc are perceived as signals.

### 2.4. Implementation Details

Previous studies [2] have shown correlations between network parameters of various protein contact networks (PCN) with the rate of folding of the proteins. GSP was used to validate these claims and check if residue signals(when visualized in frequency domain) have informative sections, which could be correlated to the rate of folding of the protein into consideration. With lower frequencies considered informative and higher frequencies as noise, absolute signal intensities in the lower eigenvalues were considered informative [14]. This informative fraction of signal is referred as Low Frequency Component (*LF C*)

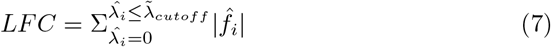

The cutoff frequency 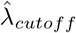 was varied from 0.01 to 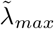 in steps of 0.01 to search for the optimum cutoff 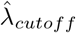 which maximizes the Pearson’s correlation between *LFC* and *ln*(*k_F_*), where *k_F_* is the rate of folding of the protein. The threshold 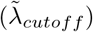 to determine the low-frequency component represents the width of the neighbourhood of co-occurrence of a property of connected residues in protein residue-network, which is correlated with folding rate. The weighted RIG distance cutoff (*r_c_*) was optimized to maximize the correlation, and 7.3 was Å fixed for further processing (Figure 2).

**Figure 2:**
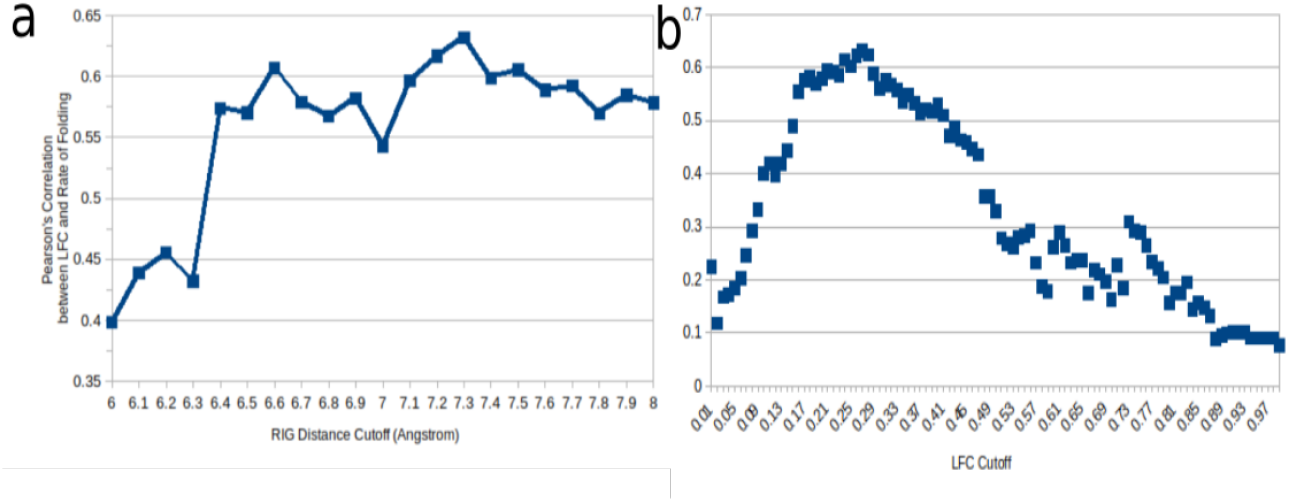
Optimizing the parameters for the Weighted RIG model with residue hydrophobicity values as signals. (a) The Euclidean distance cutoff *r_c_* was varied from 6 to 8 Å. The corresponding weighted RIG models were constructed, and for each model, the LFC cutoff was varied to obtain maximum correlation between low frequency component and folding rate. An optima were obtained at *r_c_* = 7.3 Å. (b) Scatter plot of correlation vs frequency cutoff for best performing weighted RIG model (*r_c_* = 7.3 Å). The correlation is seen to hit a maximum value of 0.63 at frequency cutoff value 0.27. The analysis is done on the 52 protein dataset, and the correlation between Log(rate of folding) and Information content in low frequencies can be observed in (Supplementary Figure 1)

### 2.5. Data Sources

In order to test our hypothesis, 52 single domain two-state folding proteins data was taken. In order to reduce model complexity, the proteins were specifically chosen to be single domain simple structures, with credible folding rate information available. Moreover, only those proteins were considered whose folding mechanism is in two-states only, without undergoing any intermediate folded structure. Such proteins were gathered from multiple sources [2, 15]. A complete list of these proteins, with their rate of folding information and protein family (*α, β* and *αβ* proteins), is given in Supplementary Table 1. Same list of 52 proteins were used for modelling *α/β* property.

## 3. Results

### 3.1. Single feature correlation model

Residue hydrophobicity is well known for its role in protein folding [16]. With hydrophobicity signal fixed, we iterated with many variations of RIG models to maximize the correlations. Kyte and Doolittle hydrophobicity scale was taken to consider signal intensities at nodes [17]. The 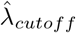 was optimized for maximum correlation between *LFC* and ln *k_F_*. As shown in figure 2b, the accumulation of the low-frequency components till certain 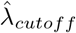 provides maximum correlation using hydrophobicity value. It indicates that when hydrophobic residues interact with each other and form a large connected group, the folding rate is high.

However, when they make smaller (below cutoff) connected groups, it has a negative impact on the folding rate. The results obtained are summarized in Table 1. The weighted RIG model was seen to outperform other RIG models, with a correlation value of 0.63.

**Table 1:** Performance Summary of different Residue Interaction Graph (RIG) models

| (Signal: Residue hydrophobicity) |  |  |  |
| --- | --- | --- | --- |
| S. No. | PCN Model | $l_{cutoff}$ | Correlation observed |
| 1 | Simple RIG | 0.27 | 0.34 |
| 2 | Unweighted RIG | 0.48 | 0.39 |
| 3 | Weighted RIG | 0.27 | 0.63 |

### 3.2. Validation against random controls

The above-mentioned results were validated against random control networks. Different network models were considered which mimic completely arbitrary proteins in order to verify the results’ sanctity. Multiple strategies or random network models were used for each protein’s weighted RIG model, a random control was designed such that the number of nodes (or residues) remains the same while the connections are changed. The edge weights were also adjusted, as being done for the weighted RIG model. Then, assuming the same hydrophobicity signal values on this weighted RC along with the same frequency cutoff, the correlations were computed. The correlation values for each random control are listed in Table 2. Each type of random control was made as close as possible to the original model as possible, in terms of the number of nodes and edges.

**Table 2:** Performance Summary for modelling folding rate when other graph is used for protein. (Signal: Residue hydrophobicity)

| Random Control Model | Correlation |
| --- | --- |
| Complete Graph | 0.07 |
| Path Graph | -0.13 |
| K-Regular Graph | 0.22 |
| Modified K-Regular Graph | -0.07 |

A detailed discussion on the methodology and implementation details pertaining to random control based validation is present in Appendix 1 (supplementary Material).

### 3.3. Prediction using multiple GSP derived features

So far, the hypothesis was only based on residue’s hydrophobicity value as a parameter for the protein folding-rate. Results in the previous sections also validate this hypothesis to a greater extent. Different signals were further used as features, and their overall effect on protein dynamics was studied. The signals chosen were both residue’s properties, like their hydrophobicity, molecular weight etc, along with the network properties like node degree, node clustering coefficient, centrality, page rank etc. To analyse the effect of different signals on the rate of folding of proteins, a linear multivariable regression model [18] was defined. Let *y* be the dependent variable, and *x*_1_, *x*_2_, … , *x_n_* are the *n* explanatory variables, a regression model tries to learn the parameters *β*_0_, *β*_1_, *β*_2_, … , *β_n_* such that 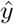 is approximated as a linear combination of the exploratory variables.

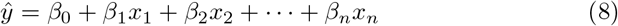

The independent predictor variables considered were critical in the regression analysis. In totality, 6 features were considered. For each of the signal, the frequency cutoff 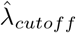 was taken to be the one which maximized the correlation between *LFC* and ln *k_F_*. The features, their corresponding 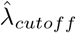 and correlation values (*ρ*) are listed in Table 3.

**Table 3:**
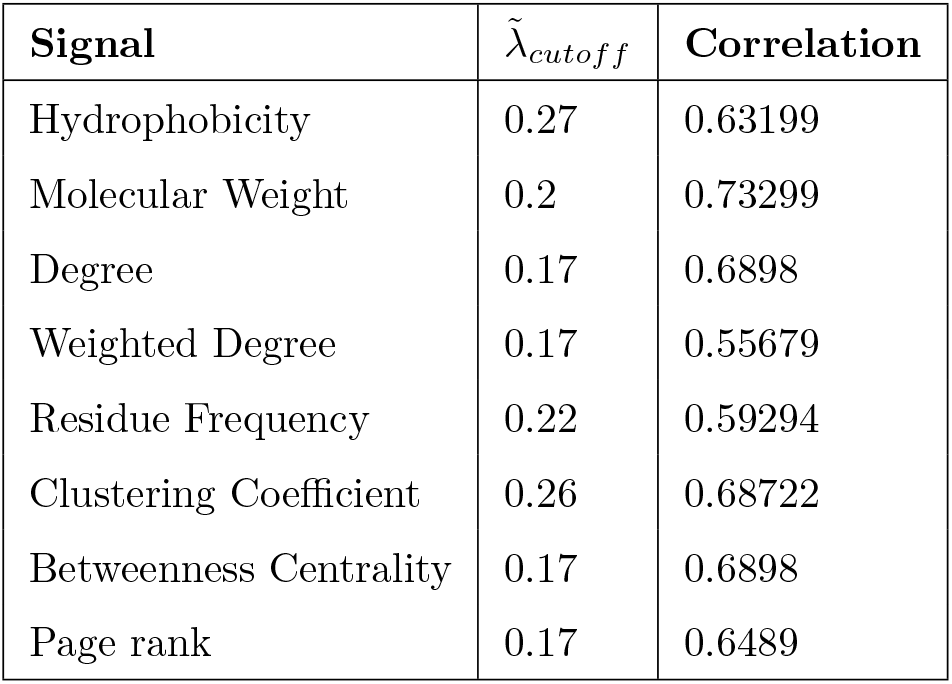
Features considered for regression model, along with their respective frequency cutoff values and correlations observed

| **Signal** | $\tilde{\lambda}_{cutoff}$ | **Correlation** |
| --- | --- | --- |
| Hydrophobicity | 0.27 | 0.63199 |
| Molecular Weight | 0.2 | 0.73299 |
| Degree | 0.17 | 0.6898 |
| Weighted Degree | 0.17 | 0.55679 |
| Residue Frequency | 0.22 | 0.59294 |
| Clustering Coefficient | 0.26 | 0.68722 |
| Betweenness Centrality | 0.17 | 0.6898 |
| Page rank | 0.17 | 0.6489 |

Once the cutoff yielding the maximum correlation was observed, the corresponding fraction of information in the cutoff section of frequency (*LF C*) was calculated across proteins. This vector of an informative fraction of the signal in the frequency domain was taken as the predictors *x_i_*. This exercise was carried out for each of the signal listed in Table 3. In this way, a feature vector for each protein was constructed. Throughout the regression analysis, the weighted RIG model was considered. The constructed feature matrix, along with folding rates, is presented in Supplementary Table 2. The proteins used in the regression model are compiled from multiple sources [15, 2]

The learned linear regression model yielded a multiple R-squared value of 0.6 (R=0.781). We performed cross-validation by randomly split of the data in 5 folds test-train samples. For each split, LASSO (Least Absolute Shrinkage and Selection Operator) based regression was used, which resulted in a 5-fold cross-validation R-square value of 0.56 (R=0.748). Our GFT based method capturing node-level properties outperforms previously reported method using holistic network properties of similarly constructed protein network models for folding rate prediction [2]. The multi-feature regression model predicted the folding rate with root-mean-square error of 1.92 and mean-absolute-error of 1.6. A comparison of other protein folding rate prediction tools done by another group is used to compare different methods [19]. As per the error reported by other methods, our approach is seen to perform better than other models [20, 21, 22] (Supplementary Table 3).

An interesting thing to observe here were the contributions of variables independently on the rate of folding[23]. Molecular weight alone contributes to roughly 20 percent of the rate of folding, as shown in Supplementary Figure 2. Other important signals are the network properties of node clustering coefficient and node degree. Residue hydrophobicity, which according to our assumptions, is seen to contribute only about 12 percent. Molecular weight is directly related to the size of amino acids, the number of atoms it has and thereby governing the number of inter-atomic attractions it can handle. [24]

### 3.4. Alpha vs Beta proteins

Tertiary structural alterations in a protein changes its resultant network model significantly. This leads to a changed eigenbasis for signal transformation. In order to examine this, we processed separately processed alpha-helices and beta-sheet proteins. Proteins which exhibited both alpha-helix and beta sheets in their tertiary structure were kept separately in a third bucket. With common signal on all the set of models, the information quotient in low frequencies were calculated. In particular, node’s clustering coefficient when used as signal resulted in a striking different *LFC* distribution along with groups, as seen in Supplementary Figure 3. The mixed class features were seen to lie between the alpha and beta classes

Deeper analysis of the alpha and beta proteins validated the efficacy of using GSP derived features in protein class identification. All the features used in the regression model earlier were calculated for these proteins. To reduce the complexity, 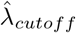 was chosen as an optimized value which was the same for each signal. Alpha, mixed and beta class proteins were labelled 1, 0 and −1 respectively, and a regression model was learned on the data. A 10-fold cross-validation based LASSO regression model resulted in an R-squared value of 0.393 (R=0.626). Further, the mixed class proteins were omitted, and the remaining proteins were used to fit another LASSO regression model. This yielded in a 10-fold cross-validated R-square value of 0.634 (R=0.796). The pairwise feature distribution (Figure 3) clearly shows a divide in GSP derived *LF C* values for different features in the two classes. The most relevant features contributing to this difference were seen to be clustering coefficient and residue molecular weight.

**Figure 3:**
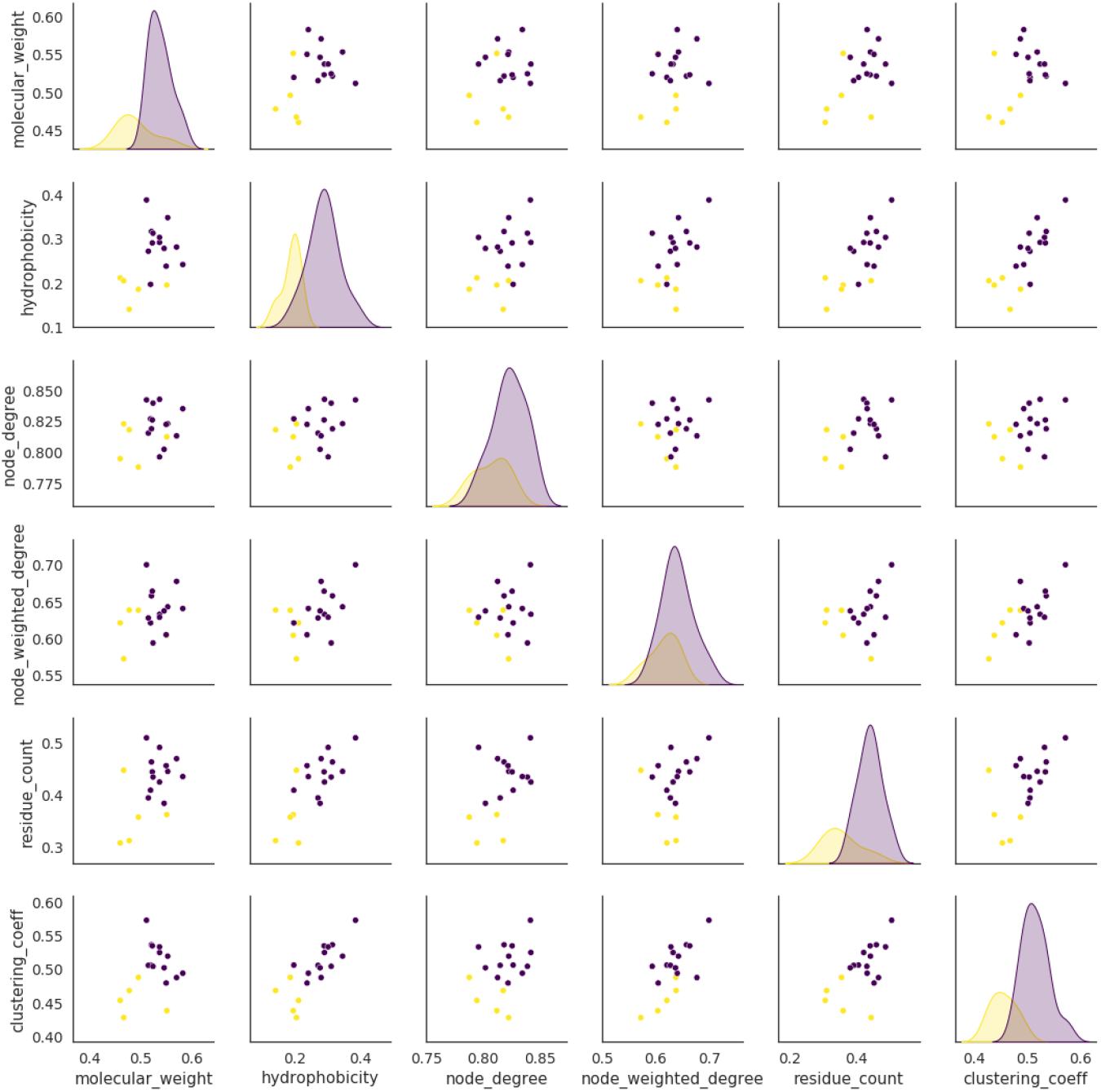
Pairwise GSP based feature distribution for alpha and beta proteins. Alpha and beta proteins are represented by yellow and blue dots respectively.

### 3.5. Transmembrane vs globular proteins

Differences in proteins’ hydrophilic and hydrophobic orientation are also observed based on the location where they are present. Proteins present in the cytosol tend to be hydrophilic in order to easily remain in a stable state. These globular proteins have hydrophilic residues towards the outer side, protecting the hydrophobic ones in the interior side of the structure. On the contrary, transmembrane proteins are a part of the lipid-rich membrane, and are thus oriented in a such a way that the hydrophobic residues are towards the external side of the structure [1]. This difference in the structural placement of hydrophobic and hydrophilic residues was assessed using the proposed frameworks. We collected a set of 237 transmembrane and 59 globular proteins and constructed their weighted RIG models. All the 8 signals were used on the models and the feature matrix was curated. This feature matrix was used to train a Logistic Regression based classification model.

The feature distribution shown in supplementary figure 4 for both the classes shows some variability in the two sets of proteins. Pairwise feature distribution indicated a greater overlap of the two classes, as seen in supplementary Figure 4. However, we utilized the power of multiple features for classification. A classification model similar to Alpha-Beta protein case study was defined for transmembrane and globular proteins. Imbalance in classes was handled by near-miss under-sampling of the over-represented group, i.e. transmembrane proteins [25]. This resulted into a 10-fold cross-validated accuracy value of 0.813, AUC score = 0.92 and F1 score of 0.822 at 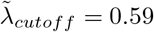, as depicted in Figure 4.

**Figure 4:**
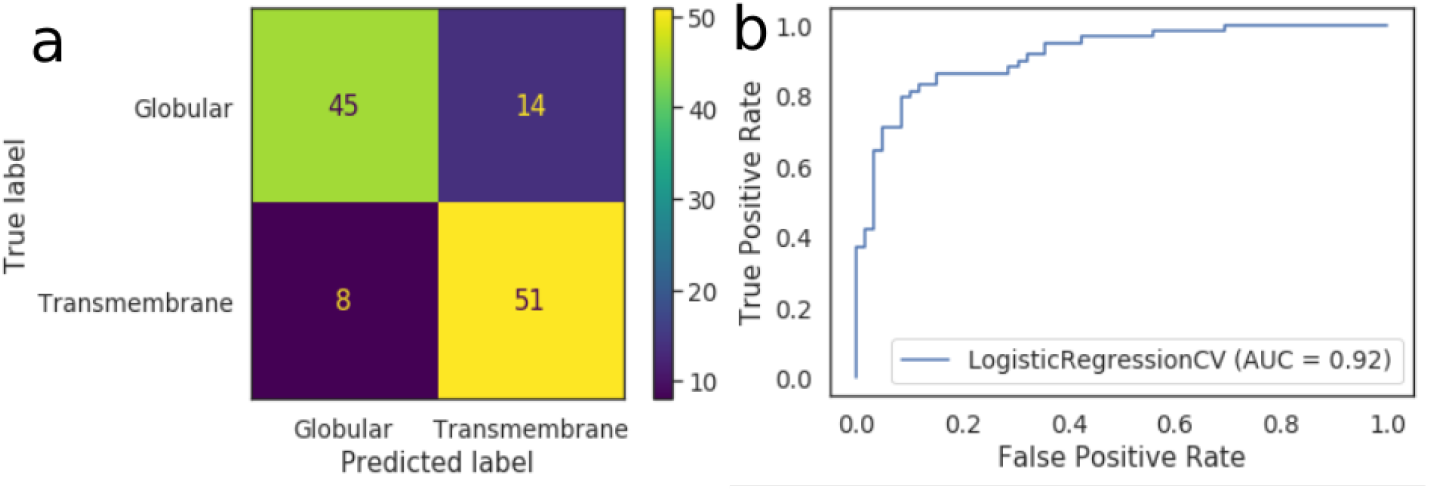
GSP derived features based Transmembrane vs Globular proteins’ classification, using 10-fold cross validation on Logistic regression. (a) Confusion Matrix of the classifier. (b) Receiver Operating Characteristic (ROC) curve for the classification

## 4. Discussion

Protein residue network can provide information to model many biophysical properties. However, most often using only graph-theoretical approach for analyzing protein residue network causes under-utilization of the information about residues’ properties. On the other hand, graph signal processing allows the use of information (or signal) of residue properties hand in hand with a graph-theoretic approach. Here, we have shown that properties of nodes based on graph theory, such as centrality, clustering coefficient can also be used together with other features of amino-acids as signal with GSP based approach.

Using a set of 52 single-domain two-state folding proteins we generated various kinds of contact networks using their known structure. The contact was decided based on Euclidean distance cutoffs and residue distance cutoffs, and residue properties are taken as signals on it. Different features of amino acids in graph Fourier domain seem to be informative about rate of folding of the protein. Such as intensity in low-frequency components while using residue molecular-weight as signal is highly informative about rate of folding of protein. It also provides insight into the modularity of protein-residue network hand in hand with residue molecular weight distribution, which affects folding rate. It hints that when heavy residues get connected with each-other in to single hub, the folding rate is high. Since we use higher weight for interaction among residues lying far from each other. It further confirms that interaction among residues with higher molecular weight and lying far from each other, leads to higher folding rate. However, if separate modules in protein fold independently of each other or heavy residues are well separated from each other, the folding rate is low. We validated our findings with a number of random controls. In this way, it is observed that using GSP, signals on a contact network can be discerned into different segments, which govern the biophysical processes involved in protein folding. A multi-variable linear regression model is also fitted, to see the contribution of various signals in folding time.

Biophysical property of protein actually emerge as a collective result of many factors. Our finding that frequency domain transformations extract informative segments with correlations with biophysical property can provide answers to unsolved puzzles regarding the distribution of amino-acids in the 3D structure of proteins. We also achieved good correlation between predicted and actual alpha/beta property (*R* > 0.76) and globularity (*R* > 0.63) of proteins using our approach of GSP and Lasso based regression model( with cross-validation). Our approach of amalgamating multiple types of signals on residue hand in hand with graph topology of proteins for machine-learning based modelling has rarely been used before. Thus our approach of extracting features using graph-Fourier transforms for down-stream machine learning approach also opens new avenues for feature extraction to predict multiple other properties of proteins. Hence in future, we would extend our approach for modelling other properties of proteins.

## Availability

All the codes for graph signal processing-based analysis of protein are available at https://github.com/divyanshusrivastava/Protein-GSP.

## conflict of Interest

There is no conflict of interest with authors

## Notes

### Competing Interest Statement

The authors have declared no competing interest.

### Summary of Updates

1. figure -2 3. changes in modelling pattern for predicting globularity

## References

[1] David L Nelson, Albert L Lehninger, and Michael M Cox. Lehninger principles of biochemistry. Macmillan, 2008.

[2] Ganesh Bagler and Somdatta Sinha. Assortative mixing in protein contact networks and protein folding kinetics. Bioinformatics, 23(14):1760–1767, 2007.

[3] Broto Chakrabarty and Nita Parekh. Naps: Network analysis of protein structures. Nucleic acids research, 44(W1):W375–W382, 2016.

[4] Wenying Yan, Jianhong Zhou, Maomin Sun, Jiajia Chen, Guang Hu, and Bairong Shen. The construction of an amino acid network for understanding protein structure and function. Amino acids, 46(6):1419–1439, 2014.

[5] András Szilágyi, Ruth Nussinov, and Péter Csermely. Allo-network drugs: extension of the allosteric drug concept to protein-protein interaction and signaling networks. Current Topics in Medicinal Chemistry, 13(1):64–77, 2013.

[6] Xuewei Jiang, Changjun Chen, and Yi Xiao. Improvements of network approach for analysis of the folding free-energy surface of peptides and proteins. Journal of computational chemistry, 31(13):2502–2509, 2010.

[7] Ganesh Bagler and Somdatta Sinha. Network properties of protein structures. Physica A: Statistical Mechanics and its Applications, 346(1–2):27–33, 2005.

[8] Helen M Berman, John Westbrook, Zukang Feng, Gary Gilliland, Tala-pady N Bhat, Helge Weissig, Ilya N Shindyalov, and Philip E Bourne. The protein data bank. Nucleic acids research, 28(1):235–242, 2000.

[9] Lesley H Greene. Protein structure networks. Briefings in functional genomics, 11(6):469–478, 2012.

[10] Nobuhiro Go and Hiroshi Taketomi. Respective roles of short-and long-range interactions in protein folding. Proceedings of the National Academy of Sciences, 75(2):559–563, 1978.

[11] Fan RK Chung and Fan Chung Graham. Spectral graph theory. Number 92. American Mathematical Soc., 1997.

[12] Aliaksei Sandryhaila and Jose MF Moura. Discrete signal processing on graphs: Frequency analysis. IEEE Transactions on Signal Processing, 62(12):3042–3054, 2014.

[13] David I Shuman, Sunil K Narang, Pascal Frossard, Antonio Ortega, and Pierre Vandergheynst. The emerging field of signal processing on graphs: Extending high-dimensional data analysis to networks and other irregular domains. IEEE signal processing magazine, 30(3):83–98, 2013.

[14] Divyanshu Srivastava and Vibhor Kumar. Graph signal processing based analysis of biological networks. PhD thesis, IIIT-D, 2018.

[15] M Michael Gromiha, A Mary Thangakani, and Samuel Selvaraj. Fold-rate: prediction of protein folding rates from amino acid sequence. Nucleic acids research, 34(suppl 2):W70–W74, 2006.

[16] H Jane Dyson, Peter E Wright, and Harold A Scheraga. The role of hydrophobic interactions in initiation and propagation of protein folding. Proceedings of the National Academy of Sciences, 103(35):13057–13061, 2006.

[17] Jack Kyte and Russell F Doolittle. A simple method for displaying the hydropathic character of a protein. Journal of molecular biology, 157(1):105–132, 1982.

[18] Sheldon M Ross. Introduction to probability and statistics for engineers and scientists. Elsevier, 2004.

[19] Catherine Ching Han Chang, Beng Ti Tey, Jiangning Song, and Ramakrishnan Nagasundara Ramanan. Towards more accurate prediction of protein folding rates: a review of the existing web-based bioinformatics approaches. Briefings in bioinformatics, 16(2):314–324, 2015.

[20] Hong-Bin Shen, Jiang-Ning Song, Kuo-Chen Chou, et al. Prediction of protein folding rates from primary sequence by fusing multiple sequential features. Journal of Biomedical Science and Engineering, 2(03):136, 2009.

[21] Chou Kuo-Chen and Shen Hong-Bin. Foldrate: A web-server for predicting protein folding rates from primary sequence. The Open Bioinformatics Journal, 3(1), 2009.

[22] Xiang Cheng, Xuan Xiao, Zhi-cheng Wu, Pu Wang, and Wei-zhong Lin. Swfoldrate: Predicting protein folding rates from amino acid sequence with sliding window method. Proteins: Structure, Function, and Bioinformatics, 81(1):140–148, 2013.

[23] Richard Harold Lindeman. Introduction to bivariate and multivariate analysis. Technical report, 1980.

[24] Alexei V Finkelstein, Natalya S Bogatyreva, and Sergiy O Garbuzynskiy. Restrictions to protein folding determined by the protein size. FEBS letters, 587(13):1884–1890, 2013.

[25] Lei Bao, Cao Juan, Jintao Li, and Yongdong Zhang. Boosted near-miss under-sampling on svm ensembles for concept detection in large-scale imbalanced datasets. Neurocomputing, 172:198–206, 2016.

